# Apparent sunk cost effect in rational agents

**DOI:** 10.1101/2021.03.26.437119

**Authors:** Torben Ott, Paul Masset, Thiago S. Gouvêa, Adam Kepecs

## Abstract

Rational decision makers aim to maximize their gains, but humans and other animals often fail to do so, exhibiting biases and distortions in their choice behavior. In a recent study of economic decisions, humans, mice, and rats have been reported to succumb to the sunk cost fallacy, making decisions based on irrecoverable past investments in detriment of expected future returns. We challenge this interpretation because it is subject to a statistical fallacy, a form of attrition bias, and the observed behavior can be explained without invoking a sunk cost-dependent mechanism. Using a computational model, we illustrate how a rational decision maker with a reward-maximizing decision strategy reproduces the reported behavioral pattern and propose an improved task design to dissociate sunk costs from fluctuations in decision valuation. Similar statistical confounds may be common in analyses of cognitive behaviors, highlighting the need to use causal statistical inference and generative models for interpretation.

## Introduction

We all strive to make good decisions that provide the maximum benefit for the least cost. However, we often succumb to a variety of cognitive biases, that is, systematic deviations from rational decisions that lead to suboptimal returns (Gilovich et al., 2002; Kahneman, 2011; Tversky and Kahneman, 1981). Understanding the behavioral and neural processes that are responsible for cognitive biases could uncover the fundamental principles behind decision making. Indeed, nonhuman animals also face decisions where the best course of action requires considering uncertainty, time, and costs. Thus, comparative studies across species can reveal insights into the biological origins of choice biases and shed light on the roots of irrational behavior (Santos and Rosati, 2015).

The sunk cost fallacy is a prominent cognitive bias, valuing an option more highly because of the resources already invested in it, instead of just considering expected future returns (Arkes and Blumer, 1985). In other words, people often stick with their poor decisions if they have already invested time, effort, or money in these decisions, even if the rational, that is, return-maximizing, behavior would be to abandon the investment and seek new opportunities. This sensitivity to sunk costs is suboptimal, thus challenging normative accounts of human decision making (Arkes and Ayton, 1999; Kahneman, 2011). However, it has been debated whether there is sufficient behavioral evidence for sunk cost-sensitive decisions in animals or, rather, if it is a uniquely human behavior (Arkes and Ayton, 1999; Magalhães and Geoffrey White, 2016). Recently, (Sweis et al., 2018a) argued that humans, mice, and rats are sensitive to sunk costs. In their tasks, the subjects had to make a sequence of decisions about how to allocate a limited time budget to gain rewards of different qualities (“web surf” task in humans (Abram et al., 2016) and “restaurant row” in rats and mice (Steiner and Redish, 2014)). Do subjects invest more time in a decision after they have already invested a lot of time? Their answer was yes: all three species seem to succumb to the sunk cost fallacy. They observed a universal behavioral pattern, one argued to be a signature of sunk cost sensitivity: the more time subjects had invested toward gaining a reward, the more likely the subjects were to keep investing until reward delivery, even when the expected future reward was the same.

Here we show that the relationship between time invested and the probability of earning a reward is subject to a statistical fallacy and arises in elementary decision models without invoking sunk costs. Therefore, the proposed behavioral signature cannot be used to infer sunk cost sensitivity. First, we provide an intuitive example of investment behavior without sunk costs to illustrate how apparent sunk cost sensitivity can arise as a consequence of a form of attrition bias, a type of selection bias that is well known in randomized controlled trials (Bell et al., 2013; Nunan et al., 2018). Next, we present a toy decision model that accounts for the choice behavior and that reproduces the reported behavioral signatures without sunk costs. Then, we provide a formal analysis of the economic decision task used by Sweis et al. (2018a) to consider the general conditions under which apparent sunk cost sensitivity can emerge. In light of our model, we also consider several additional findings presented by Sweis et al. (2018a), such as the absence of the apparent sunk cost sensitivity during offer deliberation, concluding that they do not lend further support to sunk cost sensitivity. Finally, we propose extensions to their foraging task to isolate the potential influence of sunk costs on decision behavior. Our analysis implies that direct evidence for sunk cost sensitivity in animals is still lacking, highlighting the necessity of using causal inference and generative models to interpret complex behavioral patterns (Palminteri et al., 2017).

## Results

### A rational decision maker with apparent sunk cost-sensitive behavior

Imagine a perfectly rational economist getting coffee on her way to work. One morning, her favorite coffee shop has a particularly long line. Should she still get a coffee and accept the longer wait or go next door where the line is usually shorter but where the coffee is worse? This is an investment problem in which our rational decision maker must decide whether a large investment—long waiting time in line—is worth the expected return—an excellent cup of coffee (Figure 1A).

**Figure 1.**
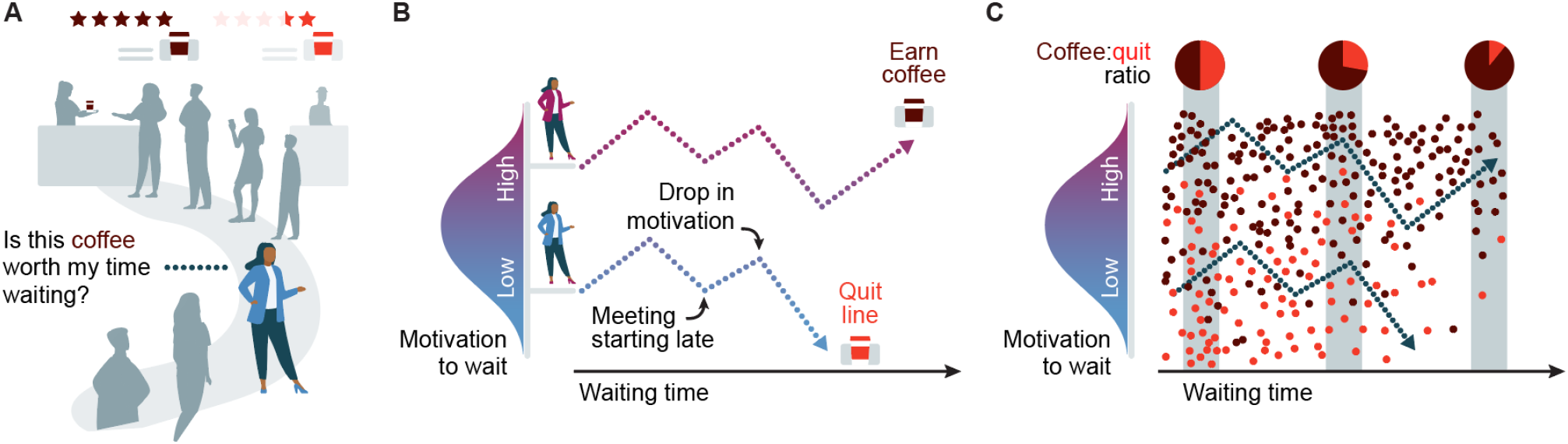
The coffee line dilemma: attrition bias produces an apparent sunk cost fallacy. **(A)** A rational decision maker deliberates whether to invest time waiting in line to get her favorite coffee. **(B)** On different days, the decision maker’s initial motivation to wait in line may be different. Her motivation while waiting in line fluctuates over time (each line corresponds to one decision to wait) due to many factors such as new information, variations in attention, or even randomly. **(C**) When following the initial decisions to wait across time (the two examples in B are shown as dashed lines), the decision maker will receive a coffee in some instances (brown dots), while in other instances she will eventually quit (red dots). However, analyzing longer waiting times (greater sunk costs) will bias the remaining observations towards higher initial motivation levels and therefore a higher likelihood of receiving a coffee. This observation bias, a form of attrition bias, leads to apparent sunk cost sensitive behavior in rational agents.

Let us first consider the coffee line across different days. On Monday morning, our economist is highly motivated to drink her favorite coffee—maybe there is a lengthy meeting ahead—so she joins the long line. However, on another day she might have decided to skip this long line. Then, once in the line, she keeps deliberating: Is the line moving fast enough? Or did the morning meeting time change at work? The motivation to keep waiting in line can fluctuate for different reasons—either because of new information or even randomly—and a significant drop will prompt our economist to quit the line and move on (Figure 1B). Importantly, without any variability in her motivation to wait, the economist would never leave the line – which is inconsistent with both our everyday experience and the behavioral patterns observed by Sweis and colleagues.

That Monday, our economist experiences a drop in motivation to wait for her coffee and decides to leave the line. Shifts in motivation will influence a rational agent, who chooses the most valuable action given the current circumstances. Despite leaving the line, our economist is a rational agent. On Tuesday morning, the line is equally long but she is even more motivated to get her favorite coffee—maybe she did not get enough sleep. Let’s suppose that she experiences identical fluctuations in motivation as the previous day, yet she stays in line until the barista finally hands her a delicious double espresso. Why did she wait so long? Did she succumb to the sunk cost fallacy?

To answer this question, we might be tempted to check if different amounts of time spent waiting (i.e., sunk costs) predicted how often our rational economist ended up getting a coffee, and to use that as a signature for sunk cost-sensitive behavior. However, the resulting behavioral pattern—longer wait times predicting a higher likelihood of receiving espresso, even when the duration of the remaining in line is the same—is confounded by varying levels of motivation. Indeed, motivation is unlikely to remain constant across days or while waiting, and even random fluctuations can trigger the decision to stop waiting. Consequently, if we examine longer waits, those will be biased toward days when her initial motivation to wait was higher to begin with (Figure 1C). In randomized controlled trials, this selection bias is known as “attrition bias”; here, a differential dropout rate of study participants (days, in our example) can introduce apparent treatment success (getting coffee, in our example) because of the “attrition” of study participants (Bell et al., 2013; Nunan et al., 2018). This statistical fallacy impedes causal inference of the factors that might influence the likelihood of getting a cup of coffee, such as sunk costs. Therefore, we cannot interpret the correlation between the time already spent waiting (i.e., sunk costs) and the likelihood of getting a coffee as evidence that sunk costs directly influence the investment decision to wait in line.

Importantly, any potential sources of variation that influence momentary motivation, from stress to attentional lapses to random fluctuations, would produce similar correlations between the waiting time and likelihood of getting the cup of coffee. In fact, even changes across days, such as increased initial motivation to enter a line for coffee as the week progresses, will lead to apparent sunk cost-sensitive behavioral patterns even in a rational decision maker who does not consider sunk costs.

In the following section, we describe a generic decision model with a rational decision maker facing the same investment decisions as in our coffee line example, here matched to the economic task and parameters used by Sweis et al. (2018a). This agent’s investment decisions are not influenced by sunk costs. However, the agent’s investment behavior shows the apparent behavioral signatures of sunk costs.

### A simple decision model produces apparent sensitivity to sunk costs without any sunk cost mechanism

First, we briefly review the behavioral design and argument for sunk costs in the study of Sweis et al. (2018a). Humans, rats, and mice were tested on how to allocate a limited time budget to gain rewards. The subjects had to first decide whether to accept or reject a time investment offer, which was the time investment required to wait for a fixed, guaranteed reward. After accepting an offer, the subjects could decide at any moment to forgo the time already invested by aborting the trial and seeking a potentially less costly (shorter time investment) reward in the next trial. Across trials, the experimenters offered different time investment durations, enabling them to compare the probability of aborting a trial for the same remaining investment time with different values of the time already invested. The authors showed that the more time the subjects had invested toward a reward, the more likely the subjects were to keep investing until reward delivery; that is, the slope of the conditional probability of staying as a function of time to reward delivery decreases with the duration of time already invested (Figure 2, Sweis et al. (2018a)). This behavioral pattern is interpreted as evidence of a sunk cost-sensitive decision mechanism.

**Figure 2:**
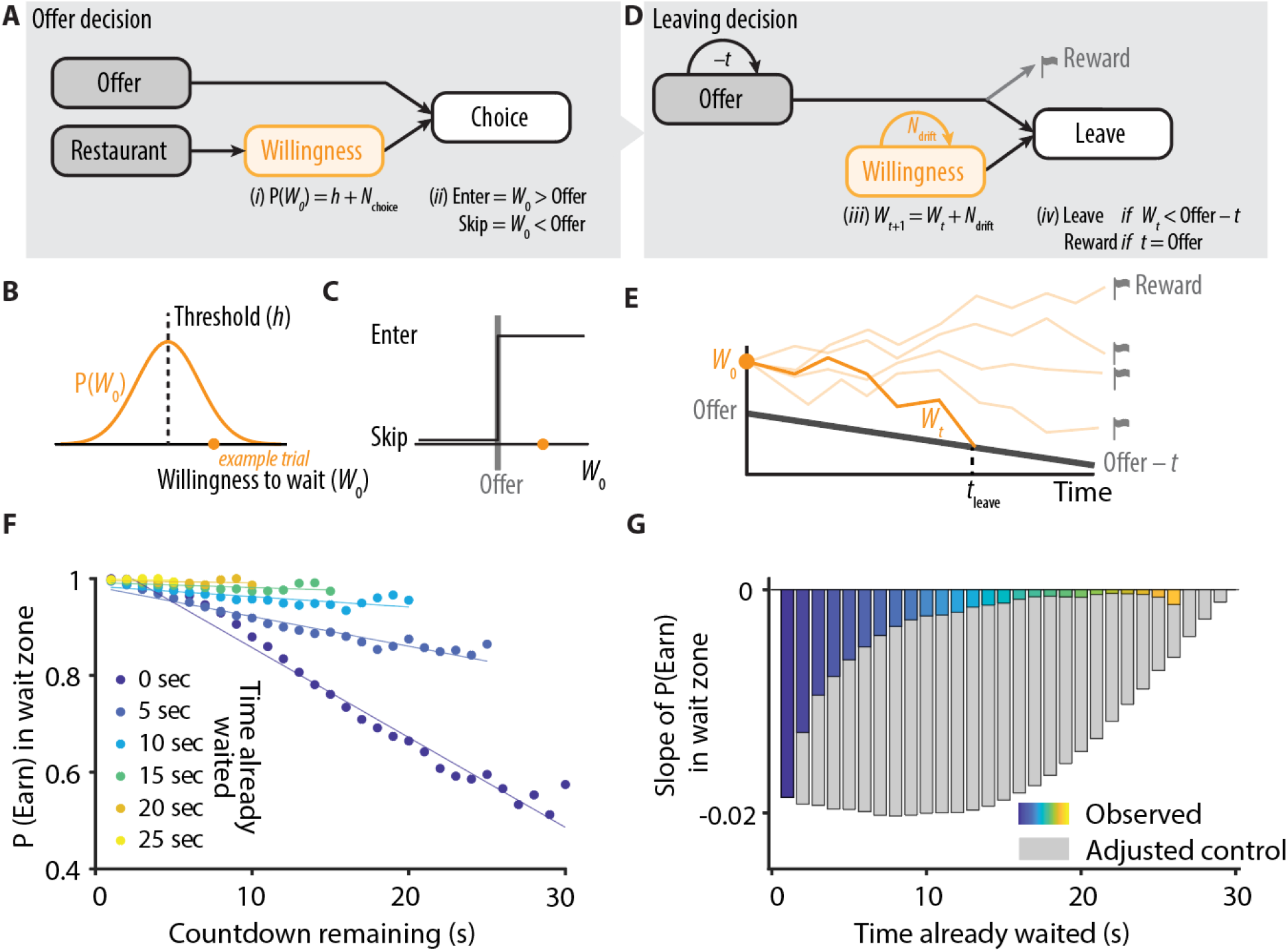
A generative model without sunk cost mechanism accounts for choice behavior and reproduces apparent sunk cost sensitivity. ***(A)** Model structure for the initial decision to accept or reject an offer. The model’s agent compares an offer value with an internal, hidden variable, the initial willingness-to-wait (W*_0_*), a noisy version of the agent’s threshold h for the current “restaurant” (equation (i)). The agent accepts an offer (and enters the wait zone) if W*_0_ > *Offer or skips an offer if W*_0_ < *Offer to proceed to the next restaurant (equation (ii)). **(B**) The initial willingness-to-wait (W*_0_*) (orange) varies across trials. When the subject is presented with an offer, W*_0_ *is sampled from a Gaussian distribution, P(W*_0_*) (defined as threshold h plus N_choice_ a zero-mean Gaussian distribution with standard deviation σ_noise_) around the threshold h. (**C**) Decision rule: The offer is accepted if the trial’s W*_0_. *is higher than the trial’s offer. (**D**) Model structure for aborting a time investment. For accepted offers, W_t_ is corrupted by noise in each second (equation (iii)) and compared with the remaining countdown Offer − t. The agent leaves the wait zone and stops investing if W_t_ < Offer − t or receives a reward if the offer time has passed, thus ending a trial (equation (iv)). (**E**) On trials in which the subject accepted the offer, W_t_ drifts during the waiting (investment) period, varying within a trial, by adding Gaussian noise N_drift_ (zero mean, standard deviation σ_drift_) in each second. Each line corresponds to one example trial with the same initial W*_0_ *and the same offer value. The drift process can either be interrupted if W_t_ drifts below the time remaining before reward delivery or if the offer time has passed and reward is delivered. The simulation was performed with h = 18 s, σ_choice_ = 5 s, σ_drift_ = 3 s, N = 1,000,000 trials. Parameters were chosen to qualitatively match the subjects’ choice behavior in the restaurant task. (**F**) Probability of earning a reward, P(Earn), after accepting an offer, that is, entering the wait zone, as a function of the remaining countdown, as well as conditioned on how long the decision agent already waited (colored lines). (**G**) Slope of the lines in (F) as a function of time already waited in the wait zone (colors as in (F)). Note that the absolute value of the slope decreases as the time already waited increases. The adjusted control corresponds to the instantaneous slope of the dark blue curve in (F) (zero time already waited) binned at the respective time already waited (gray bars)*.

Why would the subject in this task occasionally accept bad offers or stop waiting after accepting an offer? A perfect decision maker would not abort a guaranteed investment after commitment without new information or changes of reward contingencies when waiting. All species in these time investment tasks, however, showed a large variability in both their initial choices and abort behaviors. They sometimes accepted or rejected offers with the same offer time and aborted time investments, even for low offer times, that is, offer with high value (Figure S1, Sweis et al. (2018a)). In addition, the subjects sometimes abandoned an offer they had previously accepted, even though no new information about the offer was provided during the time investment, implying that there was variability in the valuation process, even within one trial. This suggests that the subjects’ investment behavior was based on a valuation mechanism, with the internal variability producing the observed choices (Cohen et al., 2007; Daw and Doya, 2006; Daw et al., 2006).

A simple toy model with elements borrowed from signal detection and drift diffusion frameworks (Bogacz et al., 2006; Gold and Shadlen, 2007; Hangya et al., 2016; Sanders et al., 2016) can account for the variability in choice behavior and produce the reported behavioral signatures that are claimed to require sunk cost sensitivity. To account for subjects’ variable choice behavior both in their initial choice (whether to accept an offer, Figure 2A) and in their investment behavior (whether to persist waiting until reward delivery, Figure 2D), we introduce an internal decision variable, *willingness-to-wait* (*W_t_*), which varies over time both across and within trials. As an internal decision variable, *W_t_* is the result of a valuation process regarding the value of waiting and, therefore, is measured in seconds. When deciding whether to accept an offer on each trial (*t* = 0), *W_0_* is sampled from a Gaussian distribution around the subject’s threshold *h*, that is, *W_0_ = h + N*_choice_ (*N*_choice_ is a zero-mean Gaussian distribution) During the time investment period (*t* > 0), *W*_t_ fluctuates following a diffusion process with noise *N*_drift_ according to *W_t+1_ = W_t_* + *N*_drift_. The initial decision to accept or reject an offer is based on the initial value of the *willingness-to-wait W*_0_ at the beginning of the trial relative to the offer *O* (in seconds) (Figure 2A–C). The decision is accepted if *W*_0_ >*O* and reject otherwise. After committing to an offer, the decision to abort an ongoing time investment is taken if the *willingness-to-wait W*_t_ drops below the remaining time before reward delivery, *O – t* (Figure 2D–E).

Using this model, we analyzed the proposed behavioral signatures of a sensitivity to sunk costs, the conditional probability of earning a reward *P*(*Earn*) as a function of time left before reward delivery *O – t* (Figure 2F,G and cf. Figure 2 in Sweis et al. (2018a)). This conditional probability was computed for the different time durations already spent waiting for the reward, that is, sunk costs *S* (Figure 2F). We observed that the slope of the curve decreased as more time had been invested (Figure 2G). Thus, the variability in *willingness-to-wait*, which is necessary to explain the variability in choice and abort behaviors, is sufficient to produce the proposed signatures of the sunk cost sensitivity without any sunk cost-sensitive decision mechanisms.

Our model, although simple, makes a few predictions. First, by construction, variability in the decision to accept or reject an offer is greatest around the subjective threshold and predicts abort decisions. The initial choices reflect a graded valuation of offer times; thus, low offer values (long-time investments) are mostly rejected, high offer values (short time investments) are mostly accepted, and there is graded variability of “accept” decisions for intermediate offers around the decision threshold (Figure 3A). Long-time investment offers that are accepted above the decision threshold (“incorrect decisions”) tend to eventually be aborted. This choice behavior is observed in all species across several studies (see Figure S1A-C, Sweis et al. (2018a), Figure 1B-E, Steiner and Redish (2014), and Figure 3, Abram et al. (2016)). Second, after accepting an offer, most decisions to abort an investment should happen early and before the remaining countdown time falls below the abort threshold, *O – t*, since the threshold moves away from the decision variable *W_t_* as time passes. This pattern of quitting behavior is observed in all species (see Figures S4 and S12, Sweis et al. (2018a)). Finally, an interesting feature of our model is that the magnitude of the apparent sunk cost effect, that is, the difference in the slopes in Figure 2F, increases with the elapsed time in a trial (Figure 2F,G). Again, this feature is observed in the data, albeit somewhat more pronounced (Figure S10C, Sweis et al. (2018a)). Note that simple additions to the model motivated by psychophysics, such as scalar timing (Gibbon, 1977), would lead to the amplification of apparent sunk cost effects for high remaining offer times.

**Figure 3:**
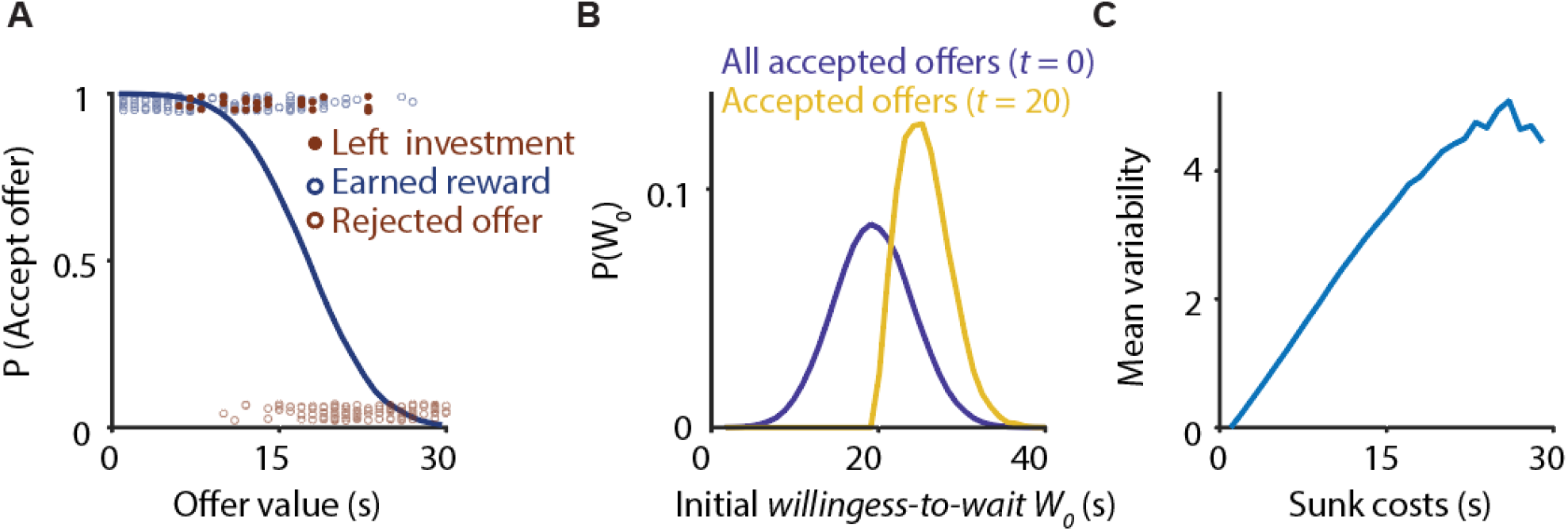
Behavioral model predictions. **(A)** Choice behavior as a function of the offer value in model implementation; see Figure 1 in Steiner and Redish (2014) and Figure 3 in Abram et al. (2016). The probability of accepting an offer decreased with increasing offer value. The data points represent 300 trials randomly sampled from the model simulation. **(B)** The distribution of initial willingness-to-wait W_0_ at the time of the offer is shifted to the right for trials in which the model subject had waited a long time (t = 20 s, i.e. sunk costs >= 20 s, yellow) compared with all accepted offer trials, i.e., trials including all waiting times (t = 0 s, i.e., sunk costs > 0 s, blue). This implies that conditioning on increasing waiting times (i.e., sunk costs) will select trials for which the drift process started on average at higher initial W_0_ values. **(C)** The mean variability N_drift_ increases with the time invested, i.e., sunk costs, S. Model parameters as in Figure 2. The observed effects were robust for a large range of tested parameters.

How does a statistical dependency between the variability in the willingness-to-wait and sunk costs arise? A behavioral analysis conditioned on how much time a subject has already waited (sunk costs, *S*) is subject to a statistical fallacy, a form of attrition bias (Bell et al., 2013; Nunan et al., 2018). Aborted trials are preferentially those that had a low initial *willingness-to-wait* values *W_0_*, because smaller random fluctuations can push them toward the abort threshold. In other words, the initial *willingness-to-wait* values *W*_0_ for all accepted trials (i.e., at *t* = 0 s) will be lower than for trials in which the subject had already waited for a longer amount of time (e.g., at *t* = 20 s) (Figure 3B). Even for the same remaining countdown time *O – t*, conditioning on longer past investments, that is, higher sunk costs *S*, will select trials with a larger initial offer *O* and, therefore, higher initial *willingness-to-wait* values *W*_0_ for the accepted offers (in which the noise *N*_choice_ has pushed *W_0_ > O*). Consequently, conditioning on higher sunk cost *S* will select more positive instances of the noise *N*_choice_, i.e. E[*N_choice_ | s_2_*] > E[*N_choice_ | s_1_*] for *s*_2_ > *s*_1_ and with *N*_choice_ | *s* representing the distribution of *N*_choice_ after conditioning on *s*. Similarly, fluctuations in *W*_t_ during waiting caused by *N*_drift_ produce a statistical dependency between *S* and the sub-selected distributions of *N*_drift_ after conditioning on *S*. Thus, the mean of noise, *N*_drift_ for a given sunk cost, *s* (i.e. E[*N*_drift_ | s]) is positively correlated with sunk cost *S* (Figure 3C). In both cases, conditioning the probability of earning a reward P(*Earn*) on sunk cost *S* will select trials with higher internal *willingness-to-wait W*_t_. This selection bias therefore cannot isolate the contribution of sunk costs to earning a reward.

### Apparent sunk cost sensitivity arises from a confounding task variable: time elapsed in a trial

In this section, we provide a formal analysis of the conditions under which apparent sunk sensitivity arises. This section generalizes the claims based on the model introduced in the previous section and can be skipped by the reader without impacting the flow of the text. We show that any fluctuation in the subject’s valuation process in determining investment decisions that either correlates with offer value or fluctuates across time is sufficient to produce apparent sunk cost-sensitive behavior.

What are the factors that determine whether a subject decides to continue to invest time in an offer or abort and move on to the test option? In the study by Sweis et al. (2018a), the analysis of the investment behavior relates to the probability of earning a reward to the time invested waiting for the reward (i.e., sunk costs) and to the time remaining to reward delivery. In each trial, a subject is presented a time investment offer *O* (we use uppercase letters for random variables and lowercase letters for values assumed by random variables). We define a subjective threshold *h* (assumed to be fixed) as the offer below which the subjects typically accept the offer. In the original study, each restaurant used a uniquely flavored food pellet as a reward (rodents) or a short video clip from a specific video category (humans), thus producing a different, but fixed, subjective threshold *h* for each restaurant or video category. Note that for simplicity and without a loss of generality, we consider a single restaurant or video category. The subjects accept an offer and start investing time if *o < h*, that is, if *o – h* < 0 (*decision rule*, the time investment offer in those trial is lower than the threshold). We define *O* = O – h* as the threshold-normalized offer at time *t = 0* to normalize the offer time across different “restaurants” (rodents) or video categories (humans). When waiting, the threshold-normalized remaining countdown time is given by *O* = [O − t] – h*. We define sunk costs *S* as the time spent investing in an offer (until reward delivery or an abort decision), that is, *s = t*. Sweis et al.’s (2018a) major finding is that the probability of earning a reward *P*(*Earn*) depends not only on the remaining countdown time *O*\* (here, *P*(*Earn*) increases with decreasing countdown time *O*\*), but also on irrecoverable sunk costs *S* (Figure 4A):

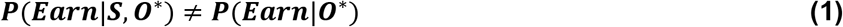

**Figure 4:**
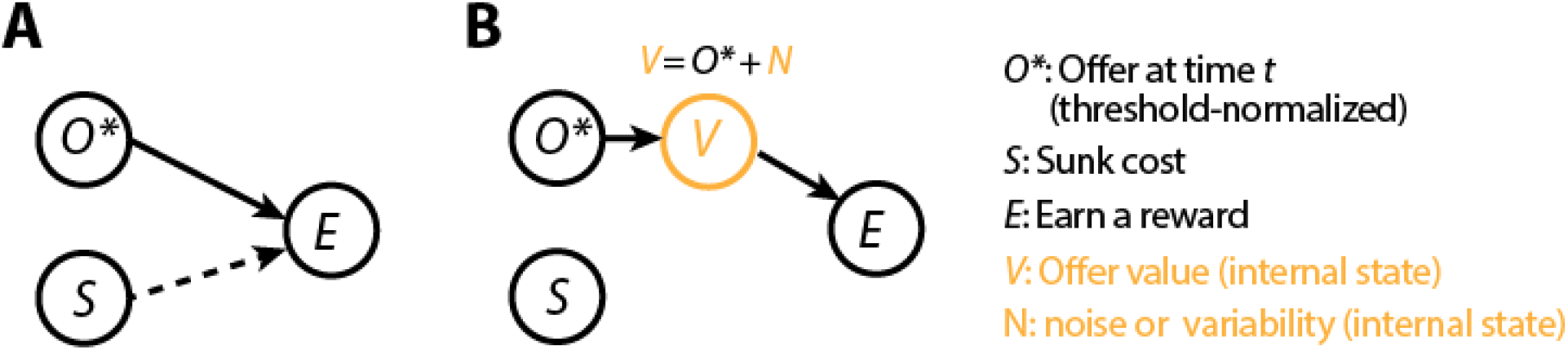
Economic decision to invest time can be driven by external variables and internal states. **(A)** Causal graphical model (a model describing a possible causal relationship between variables, Pearl, 2009) describing that the threshold-normalized offer at time t (time from investing) O* determines the investment behavior and, thus, if a reward was earned in a given trial E in the restaurant row and web surf tasks. Note that, for simplicity, we only show the most relevant model variables. The authors’ major conclusion (Sweis et al., 2018a) states that the irrecoverable time invested, sunk costs S, also influence investment decisions (dashed line). (**B**) Similar model as in (A) complemented with an additional internal state, V, describing the subjective valuation process that produces a variability in choice behavior. Observed investment behavior across humans, mice, and rats can be explained by variability in V alone but without a causal influence of sunk costs S.

Specifically, the authors show that the probability of waiting until reward delivery increases with increasing sunk costs; that is, even for the same *O*\* = *o\**, they find, for *s_2_ > s_1_*,

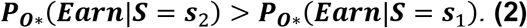

This finding (Figure 2 in Sweis et al. (2018a)) is presented as a signature of sunk cost-sensitive behavior. Interpreted causally, the decision to continue or abort the time investment is influenced by both the remaining countdown *O\** and sunk costs *S* (Figure 4A).

This interpretation does not account for the observation that accepted decisions are sometimes aborted. Indeed, there is no new information provided to subjects that would prompt them to re-evaluate their decisions. There are also no experimental interventions that would drive re-evaluation of the accepted decisions. Yet, all subjects show spontaneous aborts and in fact the entire experiment relies on these re-evaluations. We can account for both abort decisions and the original choice variability of the accept/reject decisions by assuming a noisy internal valuation process. We introduce an internal—or hidden—state of the subject: the subjective value *V* of the offer at time *t*. Because in this task value *V* refers to the value of a time investment offer, *V* can be measured in seconds. *V* can be interpreted as the subjective representation of the value of offer *O\** and, thus, expressed as *V* = – *O\** + *N*, where *N* captures the variability or noise in the subjective valuation process (in our previous toy model, *V* corresponds to the *willingness-to-wait W* with *V = W – O*). Note that high value offers of *V* correspond to short time investment offers *O*. In this hidden state decision model, an offer is accepted if V > 0 (*decision rule*) and *P*(*Earn*) is not determined by *O\** but by its internal representation *V* (here, *P*(*Earn*) increases with increasing value *V*) without the additional influence of *S* (Figure 4B):

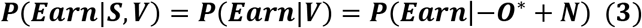

Equation 3 implies that for a fixed *O* = o\**, the probability of earning a reward is given by *P_O*_ (Earn | N)* and, thus, will statistically depend on *N*, with a higher *N* leading to higher *P(Earn)*. Importantly, if *S* and *N* are not independent, that is, *P(N | S) ≠ P(N)*, a statistical relationship between *P*_O*_(*Earn*) and *S* cannot disentangle a causal influence of either *N* or *S*. Any model in which there is a positive correlation between *N* and *S*, that is, *N* ∝ *S*, thus *V* ∝ *S* could produce qualitatively similar behavioral patterns as reported by Sweis et al. (2018a).

How could a positive correlation between the variability *N* and sunk costs *S* arise? The key feature of this task is that the sunk costs correspond to the time spent waiting for a reward, that is, *s = t*. Let us compare two distinct sunk costs amounts, *s_1_ = t_1_* and *s_2_ = t_2_* with Δ*t* = *t_2_ −t_1_* > 0. Now consider an arbitrary but fixed (threshold-normalized) remaining countdown time *o\**. Because, by definition, *o* = o – h – t*, the following conditions holds for initial offers at *t_1_* and *t_2_*

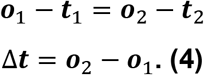

Thus, considering higher sunk costs *s_2_ > s_1_* and fixing the remaining countdown time o* will select trials with higher initial offers, *o_2_ > o_1_*. However, the valuation process *V* critically depends on the initial offer O because, by definition, an offer is accepted only when *V* > 0 at *t* = 0 (*decision rule*) for which *V(t=0) = – o*(t=0) + N(t=0) = −o + h + N(t=0)*. Crucially, even though we fixed o*, the average initial value of *V* is different when conditioning on s_1_ or s_2_. Hence, the following holds:

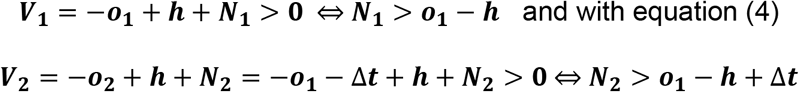

and because Δ*t* > 0, it follows

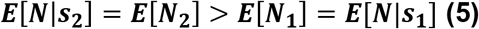

Equation (5) shows that the mean noise increases for higher investment durations s_2_ > s_1_, that is, higher sunk costs *S*. Therefore, we observe a spurious correlation *N* ∝ *S*, so the following emerges:

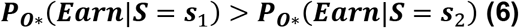

This reproduces the key behavioral observation (equation (2)) reported by Sweis et al. (2018a) (cf. toy model Figure 2, and see Figure 2, Sweis et al. (2018a)). Crucially, the statistical relation (6) holds as a consequence of the decision rule and variability in subjective valuation alone, here without a causal influence of sunk costs *S* on earning a reward *Earn*.

Moreover, any other process for which equation (5) holds will result in the statistical relation of equation (6), that is, an apparent influence of *S* on *P(Earn)*. For example, if leaving decisions while waiting for a reward are based on the momentary subjective value *V* at time *t* (e.g., leave if momentary *V* < 0), any noise in *V* will result in ***E*[*N*_2_] > *E*[*N*_1_]** because consideration in equations (4) and (5) hold not only for *t = 0* but any *t*. In other words, any temporal variability in *V* (i.e., *N*) will result in the same statistical relation, that is, an apparent sunk cost sensitivity (cf. toy model in Figure 2 and Figure 3). In addition, any positive correlation between variability *N* and offer value *O*, for example, if the variability in the valuation scales with the offer size, will also produce similar patterns, because *S* ∝ *O* (cf. equation (4)).

This analysis reveals a critical limitation in the task design for determining the sunk cost sensitivity: the valuation process and sunk costs are tightly linked through a confounding variable: time elapsed in a trial. Conditioning the probability of earning a reward on higher sunk costs *S*, that is, the time elapsed in a trial, will select instances with higher positive fluctuations *N* and, thus, a higher valuation *V*. Thus, the variability in the valuation process hinders isolating any potential influence of sunk costs *S* on earning a reward *P(Earn)* by introducing a selection bias for, on average, higher values of *N* when conditioning on an increasing *S*. Consequently, the behavioral signature proposed by Sweis et al. (2018a) (Figure 2F,G, Figure 2) does not distinguish whether the observed changes in the probability of earning a reward are because of sunk costs or the variability in the valuation process and, hence, do not provide definite evidence for sunk cost sensitivity.

### Interrupting the valuation process could dissociate sunk cost and valuation

What behavioral observations can reveal sunk cost sensitivity? Susceptibility to sunk costs is usually tested in humans by confronting subjects with two options: one “bad” choice (i.e., lower overall returns) toward a goal the subject has already invested in or an alternative “good” choice (i.e., higher overall returns) for which no prior investment has been made (Arkes and Ayton, 1999; Arkes and Blumer, 1985; Teger, 1980). Similarly, human or animal subjects could be confronted with two new choice alternatives after having already invested in one of them: In the restaurant task, we could make a novel time investment offer after subjects have already waited for a variable amount of time.

Why is it important to introduce a new offer? To isolate a sunk cost *S*, measured as the elapsed time, we need to find a way to separate it from the offer value *V* that is also correlated with elapsed time. Only an experimental manipulation can disentangle the latent correlations that naturally occur between time and value. The experimental manipulation would need to remove the arrow connecting the initial offer *O*\* and value *V* in the causal model diagram (Figure 5A). If such an experimental procedure is possible, we could then evaluate the conditional probabilities of earning a reward *P*(*Earn | S*) without a potential confound from sunk costs *S* to value *V* or noise *N*.

**Figure 5:**
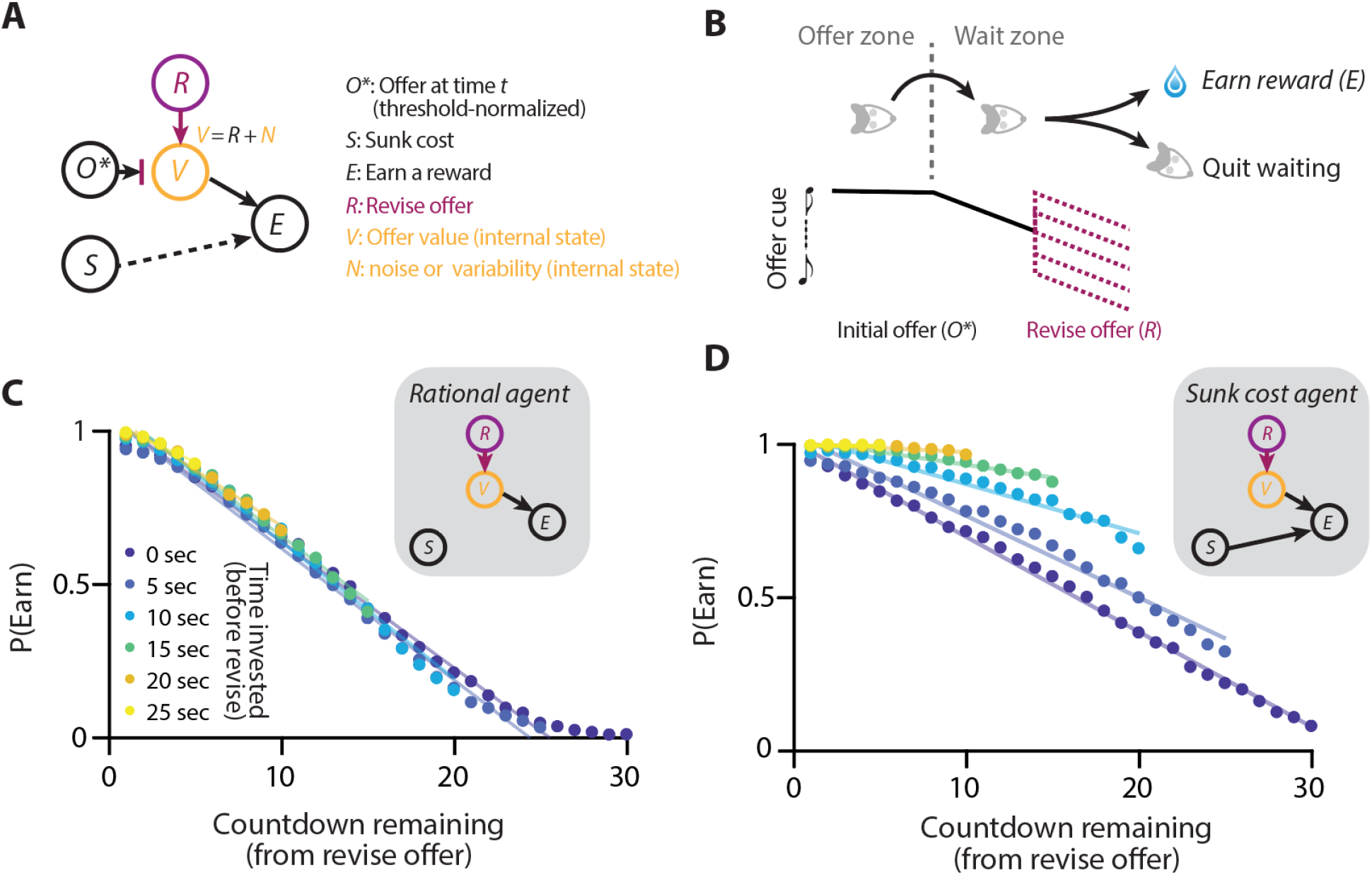
An experimental design dissociating sunk costs and noisy valuation. **(A)** Causal graphical model of the proposed behavioral manipulation. An additional revise offer (R) that is introduced randomly while waiting for the reward could remove an influence from O* to V (blocked arrow), thus allowing for a way to determine an influence from S on E (dashed arrow). **(B)** Modified restaurant task with revise offer. When waiting, a random revise offer (R) “revises” the initial offer O*. (**C**) The results of the toy model simulation of this task; the same model as in Figures 2–3 with an additional revise offer R. Timing and value of R was chosen at random between 0 s and 30 s (uniform). Probability of earning a reward P(Earn) is plotted against remaining countdown time, i.e., the value of revise offer R for different time investment values before the revise offer was shown (sunk costs, colors). As before, there was no direct influence of the sunk costs S on leaving decisions in the model. (**D**) The results of the toy model with an additional direct influence of sunk costs S on leaving decisions. Specifically, the willingness-to-wait W_t_ was increased by 1 in each second. Other model parameters are shown in Figures 2–3, except for a higher σ_drift_ = 5 s, which produces leaving decisions for the generally higher W_t_ values because of the direct sunk cost influence.

We suggest an extension to the restaurant or web surf tasks that would allow for such an experiment. If, at any moment while waiting for a reward, we “revise” the valuation process, the arrow from initial offer *O*\* to value *V* would be removed (Figure 5A). Such a revise mechanism, *R*, could be realized by randomly changing the offer value while waiting from *O*\* to *R* while making sure that *O*\* and *R* are not correlated (i.e., randomly choosing the timing and value of *R*). The current value of the offer, *V*, would then only be determined by the revise offer *R*, not by the initial offer *O*\* (Figure 5B). Behavioral signatures for sunk costs in this revise offer task are similar to the signatures previously proposed, that is, quantifying the conditional probability of earning a reward *P*(*Earn | S*), with one crucial difference. For this experiment, we do not fix the initial offer *O*\*, hence allowing for spurious correlations between sunk costs *S* and noise *N*, but we fix the momentary (threshold-normalized) remaining countdown time given by *R* = [R–t] −h* (which is defined after introducing the *r* offer *R*, i.e. *t > t*_revise_), which by its very construction is statistically not related to *O*\*. Here, comparing the different sunk costs *S* amounts to comparing trials with different time points of revise offer presentations *t*_revise_. Note that this experiment relies on the assumptions that value *V* is determined by the revise offer *R* alone and that there is no “memory” of the previous offer *O\** determining the investment decisions (and thereby preserving some degree of spurious correlation between *S* and *R*).). A possible simplification of this task could be to remove the initial offer entirely and introduce the revise offer *R* after random waiting time periods in the wait zone.

We added the random revise offer *R* to our toy model and analyzed the proposed behavioral signatures produced by the model. Indeed, when constructing conditional probabilities of earning a reward *P*(*Earn | S*) by using different sunk costs *S* at the time of the revise offer *t*_revise_ and by only using the revise offer *R* to determine the remaining countdown time, the apparent influence of the sunk costs on investment decisions was removed (Figure 5C). Next, we added an explicit sunk cost mechanism into our model by increasing the momentary willingness-to-wait *W_t_* with a fixed amount for each second of elapsed time. Analysis of the proposed behavioral signatures now reveals a sunk costs effect, as expected (Figure 5D).

## Discussion

Here we showed that a recent report arguing that humans, mice, and rats succumb to the sunk cost fallacy (Sweis et al., 2018a) is based on a value-guided decision-making task that does not allow for a dissociation between sunk costs and variability in the valuation process. An apparent sunk cost sensitivity can arise through a confounding variable in the task: the time elapsed in a trial, which reflects the sunk cost but also statistically informs the internal valuation process guiding investment decisions. Thus, although the restaurant row task provides an elegant ethological design to study valuation and economic choice (Abram et al., 2016; Steiner and Redish, 2014), it does not offer an independent measure of sunk cost sensitivity as usually understood in behavioral economics.

### Normative decision models reproduce apparent sunk cost-sensitive behavior

We presented a model that reproduces key features of the published behavioral data, but without further analyses, we do not claim that our model accounts for all aspects of the time investment behavior reported in (Sweis et al., 2018a). Nevertheless, the model clarifies how the behavioral patterns claimed to require a sunk cost mechanism can emerge by necessity from a generic decision process; hence, these signatures cannot be used as direct evidence to establish that a sunk cost mechanism is at work. In fact, numerous additional factors we did not consider could also lead or contribute to the changes in the relationship between the time invested, time remaining before the reward, and probability of obtaining a reward. Variability in motivation (Berridge, 2012; Blain et al., 2016; Steiner and Redish, 2014), perception (Lak et al., 2014; Raine and Chittka, 2007), satiety (Doya, 2008), or any other fluctuation in subjective valuation that correlates with investment durations or investment offers, including random drift across trials

Does the study by Sweis et al. (2018a) provide additional evidence for sunk cost sensitivity? An elegant feature of the restaurant row task is that there are two distinct decisions: first, whether to commit to an offer and wait (offer zone) and, second, whether to stay or quit waiting (wait zone). Only the second decision—to wait or to quit—shows apparent sunk cost sensitivity, which has been used as an argument for the specificity of this effect (Sweis et al., 2018a). However, it is unclear what factors determine the offer zone deliberation time in the first place, because the deliberation time is not related to earning a reward and there is no additional information gained while staying in the offer zone. In contrast, the waiting times after committing to an offer in the wait zone are directly related to earning a reward, and a tone signals reward proximity, continually furnishing additional information. Consequently, different decision processes likely mediate the offer and the wait zone deliberation (Sweis et al., 2018a), a proposal that is compatible with our model for time investments during waiting. For example, offer zone deliberation times (i.e., reaction time) could reflect multiple processes, including choice difficulty, attention, and motivation (Mir et al., 2011; Palmer et al., 2005; Smith et al., 2004; Zariwala et al., 2013). We did not attempt to model the complex reaction patterns observed in the restaurant row task (Sweis et al., 2018b). In fact, our account is compatible with a wide range of potential reaction time models. Indeed, any model in which wait zone deliberation times (i.e., reaction times) are not, on average, systematically related with the probability of earning a reward is compatible with the observed behavioral findings and our decision model. More generally, models of deliberation time do not constrain time investment models, nor do they provide evidence for sunk cost sensitivity of decisions.

### Alternative models for sunk cost sensitivity

How does our model relate to other proposals that explain their susceptibility to sunk costs? Statedependent valuation learning or within-trial contrast models assume that the value of an expected return is estimated relative to the current energetic or affective state (Aw et al., 2011; Pompilio et al., 2006). In these models, sunk cost sensitivity arises because the value is a decelerated function of the current energetic state, increasing the value of the same expected return the more resources are depleted. Sunk cost sensitivity in these cases also arises from an internal valuation process, similar to our model. However, both of these sunk cost models require numerous additional assumptions about how value changes with invested time and energetic state. In contrast, our model produces apparent sunk cost sensitivity through random fluctuations in the valuation process alone, accounting for the key features of the choice behavior. Therefore, our model is not an alternative to other explanations for sunk cost sensitivity; rather, it highlights how sunk cost-like behavior can arise simply because of stochasticity within a rational decision-making framework.

### Improved behavioral task design to study sunk cost sensitivity

Behavioral tasks for testing the potential effects of sunk costs need to ensure that sunk costs are not correlated with offer value or other task variables contributing to choice behavior. Uncovering the causal models underlying decisions requires behavioral manipulations or quasi-experiments to disentangle this correlation (Marinescu et al., 2018; Pearl, 2009). There are numerous behavioral designs we did not explore; for example, prompting animals at random times with offers to give up waiting for smaller rewards could probe the momentary value function underlying their abort decisions, revealing whether behavior is directly driven by sunk costs or purely by correlations between the offer value and investment size. Indeed, in the economic literature, signatures of sunk costs are often the most salient when external conditions change or are ambiguous (Camerer and Weber, 1999). These paradigms probe the idea that deviations from optimality emerge because of sunk costs, not as a consequence of random variability but of the inability to appropriately evaluate new information to maximize returns.

### Cognitive biases as statistical fallacies

Our approach could be applied to other economic decision-making scenarios. For example, in a commonly cited example of sunk costs in behavioral economics, there is the draft pick order of NBA players influenced playing time and trading strategy (Staw and Hoang, 1995). Players highest in the draft pick played more often and were traded later than players with equivalent game statistics but who were lower in the draft order, suggesting that team managers placed weight on previous, irrecoverable investments in addition to current performance. A careful analysis revealed that this sunk cost effect was greatly reduced, although still present, when accounting for latent variables, such as on-court performance or injuries. This and other examples highlight how latent variables that were unaccounted for can introduce or accentuate sunk cost–like behavioral patterns (Camerer and Weber, 1999; Kanodia et al., 1989; Mccarthy et al., 1993). In another recent study, sunk cost sensitivity was reported in two primate species trained to track a moving target with a joystick for a variable time duration (Watzek and Brosnan, 2020). Monkeys could stop and abort the trial at any time. In an analysis similar to Sweis et al. (2018a), monkeys were more likely to complete a trial and earn a reward when they had already persisted with the task for a longer period (Figure 4, Watzek and Brosnan (2020)). Again, an interpretation of this behavioral pattern will benefit from a model that considers why monkeys aborted trials at different times even for the same trial types – making this analysis susceptible to similar statistical artifacts.

Identifying decision mechanisms from behavioral observations alone is challenging because the experimenter must infer latent cognitive variables. In cognitive neuroscience and neuroeconomics, carefully designed tasks that rely on nonverbal behavioral reports have allowed researchers to relate internal variables to behavioral and neural signals, such as subjective value, motivation, attention, risk-preference, or confidence (Carrasco, 2011; Herrmann et al., 2010; Huettel et al., 2006; Kable and Glimcher, 2009; Lak et al., 2014; Lau and Glimcher, 2005; Masset et al., 2020; Ott et al., 2019; Padoa-Schioppa, 2011, 2013; Rangel et al., 2008; Sugrue et al., 2005; Vasconcelos et al., 2017). In the case of confidence, a well-known cognitive bias occurs in poor performers who are overconfident in their abilities, known as the Dunning-Kruger effect (Kruger and Dunning, 1999). This interpretation has been challenged by noting that regression to the mean would lead to similar observations of overconfidence (Krueger and Mueller, 2002; Kruger and Dunning, 2002; Nuhfer et al., 2016) and a rational Bayesian inference model largely explains the miscalibration of confidence (Jansen et al., 2021).

Our analysis also highlights the need for quantitative and causal graphical models when analyzing economic and cognitive processes (Marinescu et al., 2018; Palminteri et al., 2017). Indeed, the literature of causal statistical inference provides numerous examples for how disregarding confounder or collider variables can lead to misinterpretations (Pearl, 2009). A similar statistical fallacy is well known in clinical trials as the attrition bias, the differential dropout of study participants between treatment groups, which can lead to a misinterpretation of treatment success (Bell et al., 2013; Nunan et al., 2018). Alternatively, we can understand the present statistical fallacy as akin to the well-known Simpson’s and Lord’s paradoxes when elapsed time (i.e. sunk costs) is considered as a random variable. Here, the confounding variable is time, which both determines the sunk costs and influences the “base rates” of the variability in the valuation process when considering different sunk costs (Bickel et al., 1975; Tu et al., 2008).

In summary, we emphasize the importance of explicit models to guide the interpretation of complex behavioral processes. Counterintuitive and deceiving behavioral patterns can arise due to statistical confounds and artifact. Such statistical fallacies have been long appreciated in other fields such as econometrics and are likely to be common when investigating the behavioral signatures of cognitive processes driven by latent variables, including attention, confidence, and investment decisions (Chen and Pearl, 2012; Reynolds and Heeger, 2009; Sanders et al., 2016).

## Additional information

The Matlab code implementing the decision model will be uploaded upon publication.

## Acknowledgments

We are grateful to David Redish for the open and collegial dialogue about their study. We thank Amelia Christensen, Nathaniel Daw, Siddharth Jayakumar, Markus Meister, Camillo Padoa-Schioppa, Larry Snyder, Hao Wu, and Tony Zador for discussions and comments on earlier versions of the manuscript and G. Costa for graphical design of Figure 1. T.O. was supported by German Research Foundation (DFG) grant OT 562/1-1, P.M. was supported by the Harvard Mind Brain Behavior Interfaculty Initiative, A.K. was supported by NIH R01DA038209 and R01MH097061.

